# Wide cross-species RNA-Seq comparison reveals a highly conserved role for Ferroportins in nickel hyperaccumulation in plants

**DOI:** 10.1101/420729

**Authors:** Vanesa S. Garcia de la Torre, Clarisse Majorel-Loulergue, Dubiel A. Gonzalez, Ludivine Soubigou-Taconnat, Guillem J. Rigaill, Yohan Pillon, Louise Barreau, Sé;bastien Thomine, Bruno Fogliani, Valérie Burtet-Sarramegna, Sylvain Merlot

## Abstract

The Anthropocene epoch is associated with the spreading of metals in the environment increasing oxidative and genotoxic stress on living organisms^1,2^. Once regarded as a curiosity, plants hyperaccumulating metals are now envisioned as an opportunity to remediate metal contaminated soils. About 500 plant species adapted to metalliferous soils acquired the capacity to hyperaccumulate (>0.1% of dry weight) nickel in their shoot^3^. The phylogenetic distribution of these hyperaccumulators in 50 families suggest that this complex trait evolved multiple times independently from basic mechanisms involved in metal homeostasis. However, the exact nature of these mechanisms and whether they are shared between various lineages is not known. Here, using cross-species transcriptomic analyses in different plant families, we have identified convergent functions that may represent major nodes in the evolution of nickel hyperaccumulation. In particular, our data point out that constitutive high expression of IREG/Ferroportin transporters recurrently emerged as a mechanism involved in nickel hyperaccumulation.

---

Nickel hyperaccumulators are found worldwide as herbaceous plants, shrubs or trees growing on outcrops originating from ultramafic rocks, such as serpentine soils rich in iron, nickel, and cobalt. Although nickel hyperaccumulation is a rare and complex trait, it appeared independently in many distant plant families worldwide (**Figure 1**). In this study, we ask whether the same molecular mechanisms have been recruited convergently to reach this extreme trait. To address this question, we first compared the transcriptomes of nickel hyperaccumulators and closely related non-accumulator species to identify candidate genes involved in nickel hyperaccumulation. Then, in a second step, we established orthologous relationships to identify genes with similar functions that display differential expression in nickel hyperaccumulators from distant families.

**Figure 1.**
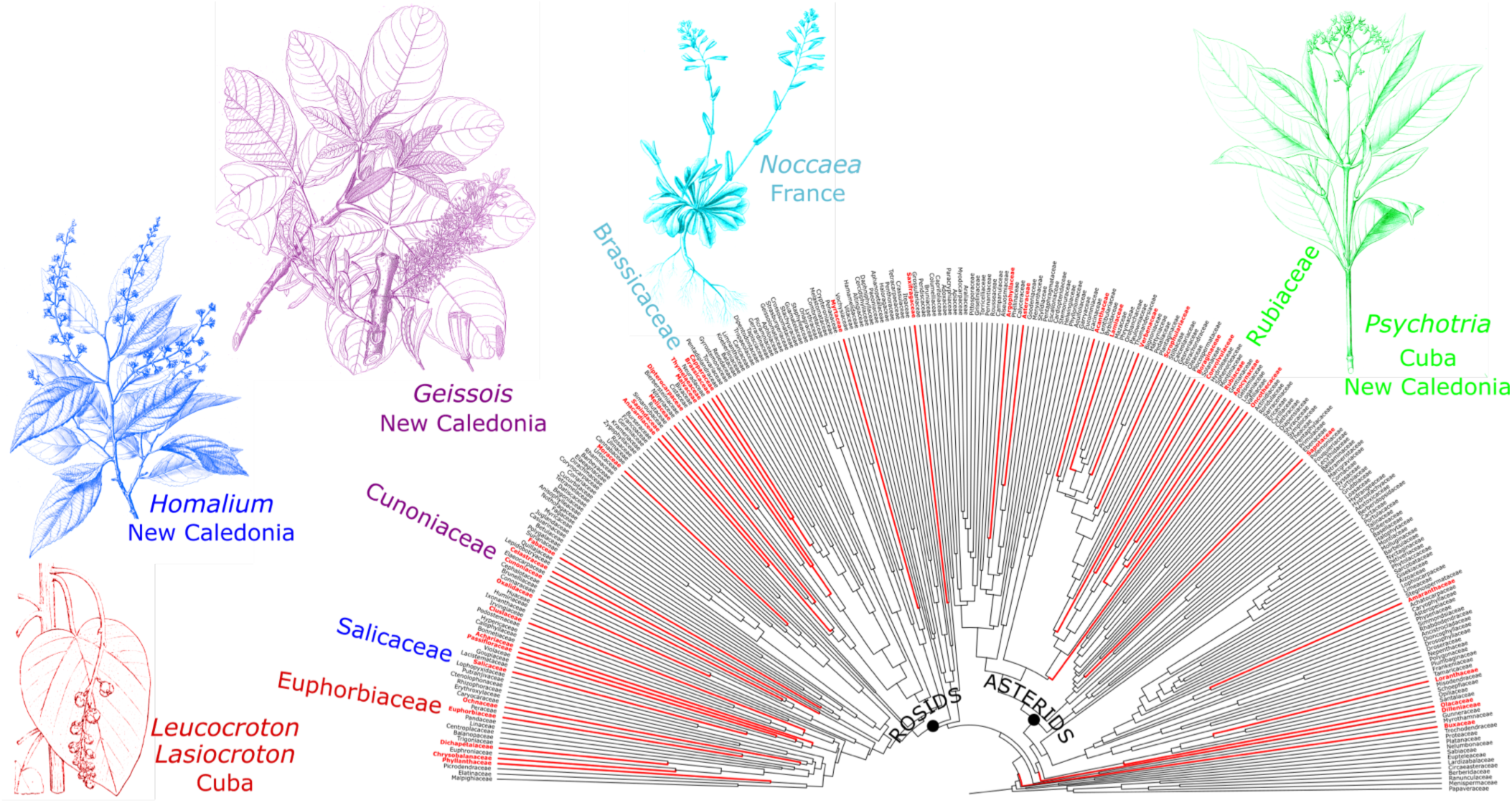
The phylogenetic distribution of nickel hyperaccumulators suggest multiple independent evolution of this complex trait. Plant families containing nickel hyperaccumulators^3^ are indicated in red on the Eudicots phylogenetic tree^4^. The drawings illustrate the plant families and genera containing nickel hyperaccumulators that we have sampled from France, New Caledonia and Cuba.

We harvested leaves of 7 nickel hyperaccumulators from 5 genera (i.e, *Noccaea, Psychotria, Geissois, Homalium, Leucocroton*) corresponding to distant plant families in their natural environment in France, New Caledonia and Cuba (**Figure 1**, **Table 1)**. For each hyperaccumulator, we collected leaves of a closely related species or ecotype not accumulating nickel in the same geographic area. To identify genes differentially expressed between closely related nickel hyperaccumulator and non-accumulator species, we performed RNA sequencing (RNA-Seq) using Illumina paired-end technology (**Supplementary Data 2**). In absence of genomic sequence references for most of these species, we *de novo* assembled the short Illumina reads into the sequence of expressed genes in the 13 selected species. These newly generated transcriptomes contain between 41,843 and 87,243 contigs for a total assembly size ranging from 35 Mbp to 49 Mbp **(Table 1, Supplementary Data 2**). Subsequent annotation of these transcriptomes using Blastx interrogation of Viridiplantae proteins database revealed significant homologies for 56 % of the contigs on average (E-value ≤10E-6). These high-quality reference transcriptomes constitute a unique and comprehensive resource for molecular studies on plant groups of high interest.

**Table 1.** *De novo* assembled transcriptomes of nickel hyperaccumulators and related non-nickel accumulator species.

| Family/ Species | Trait* | Origin | Nbr. contigs | Assembly size (Mbp) | N50 (bp) |
| --- | --- | --- | --- | --- | --- |
| **Brassicaceae - *Noccaea*** |  |  |  |  |  |
| *Noccaea caerulescens* subsp. *firmi*ensis (Ncfi) | NiH | France | 41,843 | 43.6 | 1,735 |
| *N. caerulescens* Viviez (Ncviv) | ZnH | France | nd* | nd* | nd* |
| *N. montana* (Nmon) | NA | France | 87,243 | 48.8 | 692 |
| **Rubiaceae - *Psychotria*** |  |  |  |  |  |
| *Psychotria gabriellae* (Pgab) | NiH | New Caledonia | 60,899 | 45.5 | 1,140 |
| *P. semperflorens* (Psem) | NA | New Caledonia | 66,755 | 49.8 | 1,181 |
| *P. grandis* (Pgra) | NiH | Cuba | 45,143 | 36.8 | 1,344 |
| *P. costivenia* (Pcos) | NiH | Cuba | 46,451 | 35.1 | 1,240 |
| *P. clementis* (Pcle) | NA | Cuba | 56,754 | 44.0 | 1,245 |
| **Cunoniaceae - *Geissois*** |  |  |  |  |  |
| *Geissois pruinosa* (Gpru) | NiH | New Caledonia | 54,969 | 42.1 | 1,243 |
| *G. racemosa* (Grac) | NA | New Caledonia | 58,386 | 43.7 | 1,021 |
| **Salicaceae - *Homalium*** |  |  |  |  |  |
| *Homalium kanaliense* (Hkan) | NiH | New Caledonia | 52,634 | 40.6 | 1,259 |
| *H. betulifolium* (Hbet) | NA | New Caledonia | 49,962 | 39.0 | 1,294 |
| **Euphorbiaceae – *Adelieae* tribe** |  |  |  |  |  |
| *Leucocroton havanensis* (Lhav) | NiH | Cuba | 58,990 | 44.4 | 1,206 |
| *Lasiocroton microphyllus* (Lmic) | NA | Cuba | 52,421 | 39.8 | 1,244 |

\* NiH (Ni hyperaccumulator), ZnH (Zn hyperaccumulator), NA (non-accumulator), nd (nondetermined)

The reads from each sample were then mapped to the nickel hyperaccumulator reference transcriptome to identify differentially expressed (DE) genes in 8 pairs of species, each containing a hyperaccumulator and a related non-nickel accumulator. The fraction of DE genes ranged from 2 % to 29 % (Fold Change ≥ 2, *FDR* ≤ 0.05), depending on the pair of species considered (**Figure 2a, Supplementary Figure 2 and Supplementary Data 4-9**). Swapping to the transcriptome of the non-accumulators as reference had only a marginal influence on the results (**Supplementary Figure 3**).

**Figure 2.**
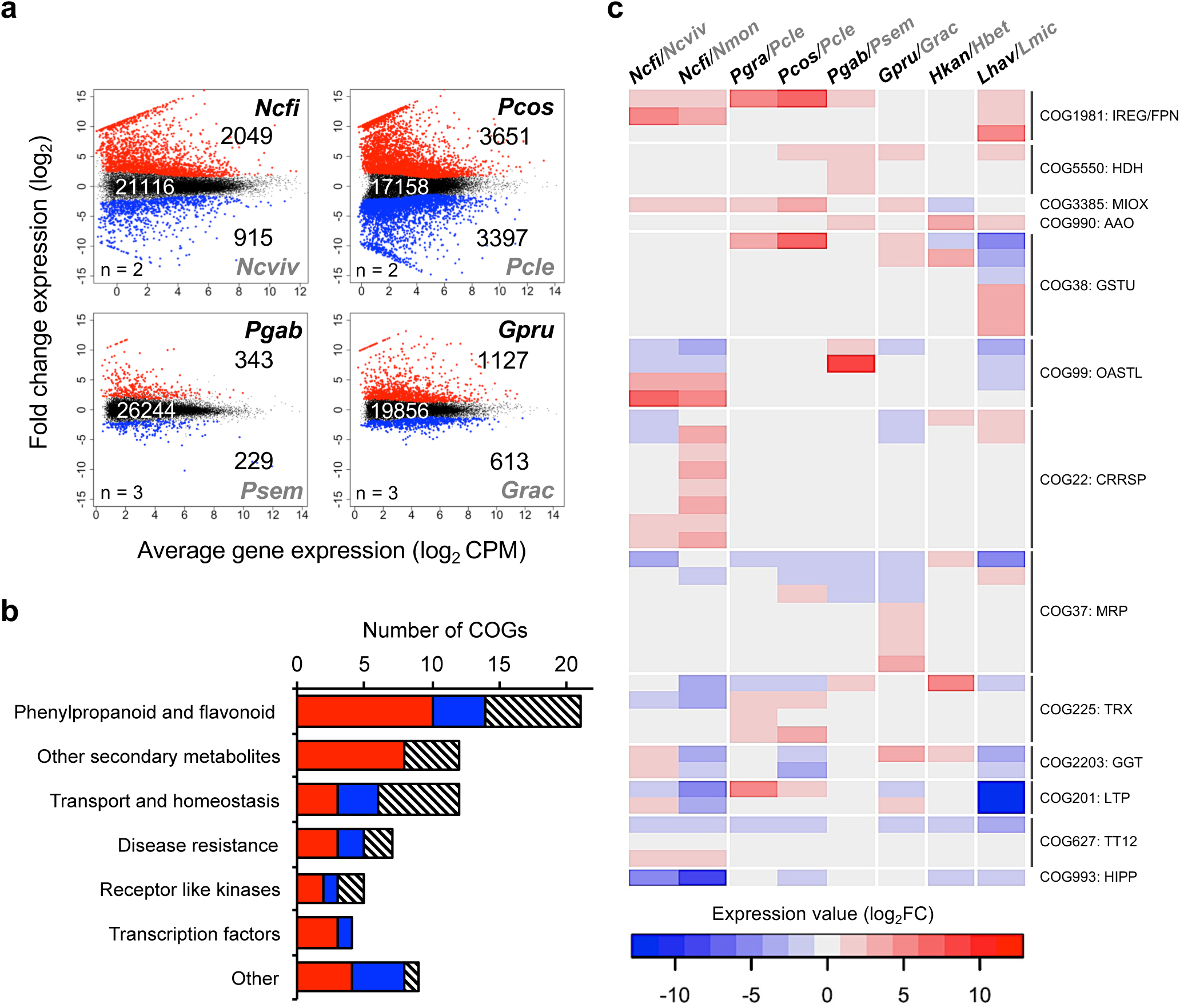
Cross-family comparison reveals orthologous genes convergently associated with nickel hyperaccumulation. (a) Cross-species comparative transcriptomic reveals Differential Expressed (DE) genes between nickel hyperaccumulator species (black) and related non-nickel accumulators (grey). Genes (contigs) are plotted according to their mean level of expression (x-axis) and their differential expression (y-axis) in the pair of species. The numbers of significant DE (red and blue dots) and non DE genes are indicated (log2FC>1, *FDR<*0.05).(b) Distribution of the 71 selected Cluster of Orthologous Groups (COGs) associated with nickel hyperaccumulation according to their predicted function and their level of expression in hyperaccumulators: Red represents COGs containing DE genes more expressed in nickel hyperaccumulators belonging to at least 3 distinct plant families, the blue color for COGs containing DE genes less expressed in nickel hyperaccumulators, and hatched for COGs containing DE genes both more and less expressed in nickel hyperaccumulators. (c) Heat-map of COGs containing genes related to transport and homeostasis function that are differentially expressed in nickel hyperaccumulators compared to related non-accumulators. Abbreviations: IREG/FPN (Iron Regulated/ Ferroportin), HDH (Histidinol Dehydrogenase), MIOX (Myo-Inositol Oxygenase), AAO (Ascorbate Oxidase), GSTU (Glutathione-s-transferase Tau), OASTL (Cysteine Synthase), CRRSP (Cysteine-rich Repeat Secretory Protein), MRP (Multidrug Resistance Associated Protein/ABCC), TRX (Thioredoxin-H-type Protein), GGT (Gamma-Glutamyltransferase), LTP (Lipid Transfer Protein), TT12 (Mate Transparent Testa12), HIPP (Heavy Metal-associated Isoprenylated Plant Protein).

**Figure 3.**
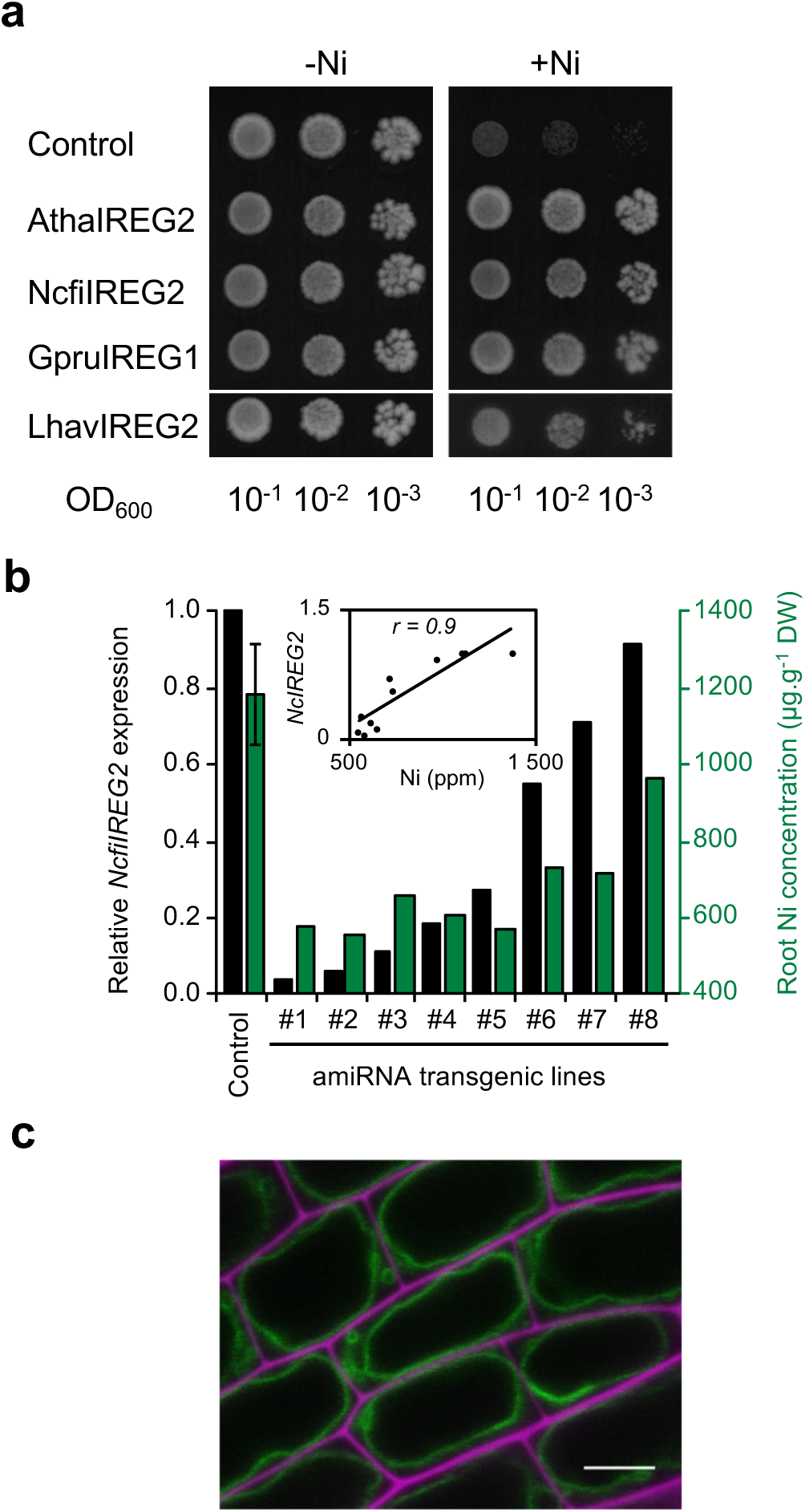
IREG/Ferroportin transporters have a conserved role in nickel accumulation. (a) Expression of plant IREG/Ferroportin transporters increases nickel tolerance in yeast. Yeast cells expressing IREG/Ferroportin transporters cloned from *A. thaliana, N. caerulescens, G. pruinosa and L. havanensis* were plated at different dilutions on a medium containing a toxic concentration of nickel for the control line (transformed with pDR195 vector). (b) Silencing of the IREG/Ferroportin transporter *IREG2* in roots of the nickel hyperaccumulator *N. caerulescens subsp. firmiensis* reduces nickel accumulation. The expression of *NcIREG2* was quantified by RT-qPCR (black bars) in 8 amiRNA transgenic lines and control lines transformed with the pK7WG2D vector (n=3) growing in presence of 37.5 μM NiCl_2_ for 4 weeks. *NcIREG2* expression was corrected using *Nc6PGDC* as reference gene and normalized to 1 for the expression in control lines. Nickel concentration was measured in parallel in the roots of the same lines by MP-AES (green bars). The insert presents the Pearson’s linear correlation between *NcfiIREG2* expression and nickel accumulation (*P-value* < 0.001). (c) NcfiIREG2 localizes on the vacuolar membrane in *N. caerulescens* cells. Confocal picture of a transgenic line expressing NcfiIREG2-GFP (green) in root cells. Cell wall was stained with propidium iodide (magenta). The scale bar correspond to 5µm.

To determine if deregulated molecular and metabolic pathways are shared by nickel hyperaccumulators from distant plant families, it is necessary to compare these transcriptomes together with a functional emphasis. To this aim, we classified the gene products into Clusters of Orthologous Groups (COG). Combining the 13 reference transcriptomes, 443,400 contigs (66.5% of total) were assigned to 46,458 COGs (**Supplementary Data 3**).

To target genes involved in nickel hyperaccumulation, we looked for COGs containing differentially expressed genes in at least 3 plant families. This analysis revealed 71 groups, of which 33 COGs included orthologous genes more expressed in nickel hyperaccumulators, 15 COGs included orthologous genes less expressed and 22 COGs were heterogeneous with some orthologs more expressed and some orthologs less expressed (**Supplementary Data 10**).

The functional annotation of these COGs **(Figure 2b)** indicated that the most represented category corresponds to genes involved in the biosynthesis of secondary metabolites, more specifically phenylpropanoids and flavonoids molecules **(Supplementary Figure 4)**. The analysis of gene expression in these pathways predicts an increased synthesis of flavonoids in leaves of nickel hyperaccumulators. This suggests an unsuspected role for flavonoids in nickel hyperaccumulation. These molecules have been shown to act as antioxidants or directly as metal ligands, including nickel, *in vitro*^5,6^. Furthermore, it has been recently shown that flavonoids accumulate in an *Arabidopsis halleri* population hyperaccumulating cadmium^7^. Further metabolomics, speciation and genetics studies will be required to establish the importance of flavonoids in nickel hyperaccumulation.

Previous studies have highlighted the important role of histidine in nickel hyperaccumulation in Brassicaceae^8^. Our data reveal that orthologs of histidinol dehydrogenase (COG5550), catalyzing the last step of histidine synthesis, are more expressed in nickel hyperaccumulators from Rubiaceae, Cunoniaceae and Euphorbiaceae families supporting the role of histidine in nickel hyperaccumulation (**Figure 2c)**. Interestingly, in the hyperaccumulator *Alyssum lesbiacum* of the Brassicaceae family, the first step of histidine biosynthesis (ATP-PRT) is upregulated^9^. This result suggests that distinct steps of the histidine biosynthetic pathway are amplified converging to increased histidine synthesis in nickel hyperaccumulators from different families.

Our cross-species comparison points to IREG/Ferroportin transporters (COG1981) as the most robust up-regulated function among distantly related nickel hyperaccumulators (**Figure 2c, Supplementary Data 10**). IREG/Ferroportin (SLC40) are efflux transporters found in different kingdoms and displaying broad specificity for divalent metal ions^10,11^. In vertebrates, the main function of IREG/Ferroportin is to release iron from cells^12–14^. In plants, IREG/Ferroportin transporters have been associated with the regulation of metal homeostasis including iron, nickel, cobalt and aluminum^15–19^. Genes encoding IREG/Ferroportin transporters are significantly more expressed (from 4 to 800-fold increase) in nickel hyperaccumulators from the Brassicaceae, Rubiaceae and Euphorbiaceae families compared to their related non-accumulator species. In Salicaceae and Cunoniaceae species from New Caledonia, *IREG/Ferroportin* genes are highly expressed in both hyperaccumulators and non-accumulators (**Supplementary Figure 5a**). However, the unique IREG/Ferroportin gene (COG1981) detected in the hyperaccumulator *Geissois pruinosa* is 60-times more expressed than its orthologue in the non-accumulator Cunoniaceae species *Cunonia capensis* from South Africa (**Supplementary Figure 5b**). Therefore, the high expression of *IREG*/*Ferroportin* genes in non-accumulator species endemic from New Caledonia might be a genetic footprint of the recent colonization (∼35 MYA) of this island then probably fully covered by an ultramafic rock layer^20,21^.

To provide functional evidence for the role of plant IREG/Ferroportin in nickel hyperaccumulation, we expressed IREG/Ferroportin orthologs cloned from 3 distant nickel hyperaccumulator species in yeast (**Figure 3a**). The expression of these transporters increases yeast resistance to nickel, which is consistent with a conserved activity of plant IREG/Ferroportin as nickel exporters. Then we investigated the biological relevance of IREG/Ferroportin in nickel hyperaccumulation *in planta*. We used *Rhizobium rhizogenes* transformation to silence the expression of *NcIREG2* in the roots of the nickel hyperaccumulator *N. caerulescens* subsp. *firmiensis* using artificial miRNA technology. We chose to target *NcIREG2* which shows strong differential expression in leaves and roots associated with nickel hyperaccumulation in *Noccaea caerulescens* (this study and ^22^). We generated 8 independent transgenic lines displaying different degrees of *NcIREG2* silencing (**Figure 3b**). Elemental analysis of these transgenic lines revealed a decrease of nickel accumulation in roots. Moreover, the amplitude of nickel accumulation strongly correlates with *NcIREG2* expression (Pearson’s *r* = 0.9, p-value = 5.3E-5). To understand the cellular basis for NcIREG2 mediated nickel accumulation, we expressed a GFP tagged version of this transporter in roots. Confocal imaging of transgenic roots shows that NcIREG2-GFP localizes on the membrane of the vacuole. Together, our results provide genetic evidence that NcIREG2 contributes to nickel hyperaccumulation by driving nickel sequestration in vacuoles.

The development of RNA-Seq technologies has opened the possibility to study non-model species at the molecular level. Yet, comparative biology has not fully benefited from this revolution because of the difficulty to quantitatively compare transcriptomes from distant species. In this study, we have used a combination of cross-species comparative transcriptomics analysis and COG annotation to identify genes associated with nickel hyperaccumulation in a wide diversity of plant families. This analysis revealed a limited number of candidate gene functions corresponding to convergent mechanisms involved in nickel hyperaccumulation including flavonoid and histidine biosynthesis, and nickel transport. Strikingly, constitutive high expression of IREG/Ferroportin in leaves has been recurrently recruited as a convergent mechanism for nickel hyperaccumulation. Our functional analysis provides evidence that these transporters account for the exceptional ability of hyperaccumulators to store nickel in vacuoles. This shows that our transcriptomic approach predicts important actors in nickel hyperaccumulation. This study further highlights other candidate genes as target for future functional analyses. Moreover, it provides a framework to identify key genes from root transcriptomes that are involved in the efficient uptake and translocation of nickel to the leaves. The identification of all molecular steps from uptake in roots to sequestration in leaves is necessary to fully understand this complex trait. In the context of sustainable development, nickel hyperaccumulators are now viewed as crops to extract and recycle metals from large areas of metalliferous soils^23,24^. As for other crops, we foresee that this molecular knowledge could become instrumental for marker-assisted selection of cultivars or molecular monitoring of agricultural practices to improve nickel phytoextraction.

## METHODS

Methods, including statements of data availability and associated accession codes and references, are available in the online version of the paper.

*Note: Any Supplementary Information is available in the online version of the paper*

## ACKNOWLEDGMENTS

We thank professor Rosalina Berazaín Iturralde (UNAH, Cuba) for invaluable information on Cuban flora, Louis-Charles Brinon (IAC, New Caledonia) for sample collection in New Caledonia, Véronique Brunaud and Marie-Laure Martin-Magniette (IPS2, France) for guidance on *de novo* assembly and biostatistical analysis, and Mark G. M. Aarts (WUR, Netherlands) for the *Noccaea* root transformation protocol.

This work was supported by Grants ANR-13-ADAP-0004 (SM, BF, VBS) and CNRS Defi Enviromics Gene-4-Chem to SM, a SCAC fellowship from the French Embassy in Cuba to DAGand SM and an ATIGE grant from Génopole to GJR. The I2BC and POPS platform benefit from the support of the LabEx Saclay Plant Sciences-SPS (ANR-10-LABX-0040-SPS). This work has benefited from the core facilities of Imagerie-Gif, a member of Infrastructures en Biologie Santé et Agronomie (IBiSA), supported by France BioImaging Grant ANR-10INBS-04-01 and the Saclay Plant Science Labex Grant ANR-11-IDEX-0003-0. We thank the South Province of New Caledonia and the Prefecture of Aveyron for plant collection authorizations.

## AUTHOR CONTRIBUTIONS

BF, VBS and SM designed the project; VSG, CML, DAG, BF, VBS and SM collected plant samples; VSG, CML, DAG, LB performed experiments; LST supervised RNA-Seq sequencing; VSG, GJR, YP, ST and SM analyzed the data; VSG, ST and SM wrote the manuscript; all authors commented and approved the content of the manuscript.

## COMPETING FINANTIAL INTERESTS

The authors declare no competing financial interests.

## ONLINE METHODS

### Plant material and sample collection

Leaves of nickel hyperaccumulator plants belonging to 5 distinct plant families^25–28^, Brassicaceae [*Noccaea caerulescens subsp. firmensis* (*Ncfi*)], Cunoniaceae [*Geissois pruinosa (Gpru)*], Euphorbiaceae [*Leucocroton havanensis (Lhav)*], Rubiaceae [*P. costivenia (Pcos), P. gabriellae (Pgab), Psychotria grandis (Pgra)*], Salicaceae [*Homalium kanaliense (Hkan)*], and of related non-accumulator species, Brassicaceae [*N. caerulescens* “Viviez” (*Ncviv*), *N. montana* (*Nmon*)], Cunoniaceae [*G. racemosa (Grac)*], Euphorbiaceae [*Lasiocroton microphyllus (Lmic)*], Rubiaceae [*P. clementis* (*Pcle*), *P. semperflorens (Psem)*], Salicaceae [*H. betulifolium (Hbet)*], were collected from individual plants growing in their natural environment in France, New Caledonia and Cuba (**Supplementary Data 1**). We complied with local regulation for the access to these genetic resources. Each sample was localized by GPS. For each sample, a fraction of leaves was washed with water and dried for elemental analysis, and the other fraction fixed on site with liquid N2 (*Gpru, Grac, Pgab, Psem*) or with RNAlater (Sigma Aldrich) and stored at 4°C (*Ncfi, Ncviv, Nmon, Lhav, Lmic, Pcos, Pgra, Pcle, Hkan, Hbet*). RNAlater was removed in the laboratory and leaves were immediately stored at −80°C before RNA extraction.

Additionally, to generate reference transcriptomes, *Ncfi* and *Nmon* were grown in hydroponic condition using a modified Hoagland’s solution^29^ containing 20 μM Fe-HBED (Van Iperen International) and 37.5 μM NiCl_2_, in a climatic chamber (9 h light, 150 μE m^-2^ s^-1^, 21 ºC/17 ºC day/night; 70 % humidity) for 7 weeks. *L. havanensis* seeds were cultured *in vitro* on Murashige & Skoog Agar medium supplemented with 3.2 mM NiSO_4_, as described^30^.

A specimen of *Cunonia capensis* (*Ccap*) was obtained from a specialized plant nursery (Ets. Railhet, France) and grown in a green-house on coconut fiber supplemented with fertilizer.

### RNA sequencing

Total RNA from leaves were extracted with RNeasy Plant Mini kit

(Qiagen) for *Noccaea* species, Qiagen hybrid method for woody plants^31^ for *Psychotria* species from Cuba, CTAB-PVP method ^31^ for *Geissois* and *Psychotria* species from New Caledonia, and TRI Reagent (Sigma-Aldrich) for *Leucocroton, Lasiocroton* and *Homalium* species. DNA was removed from all RNA samples by RNeasy Plant Mini kit on-column DNase I treatment.

RNA quality control, preparation of cDNA libraries, sequencing and raw reads processing were performed by the POPS transcriptomic platform (IPS2, Orsay, France). Libraries were prepared from 1 µg of total RNA using TruSeq Stranded mRNA kit (Illumina) and sequenced with an Illumina HiSeq2000 sequencing system in 100bp paired-end mode. Libraries meant to be directly compared were multiplexed and sequenced in a single run. Adaptors and low-quality pair-end sequences were removed from the raw reads and the ribosomal RNA was filtered using the SortMeRNA algorithm^32^. We obtained between 27 and 106 million reads per libraries (**Supplementary Data 2**).

### *De novo* transcriptome assembly and annotation

The transcriptome sequences of *Ncfi, Nmon, Pgab, Psem, Pgra, Pcos, Pcle, Gpru, Grac, Hkan, Hbet, Lhav* and *Lmic* were obtained independently by *de novo* assembly of paired-end reads using CLC Genomics Workbench v9 software (Qiagen). A single library per species was used to minimize genetic variability. Assembly parameters were set as default, except similarity (0.95), length fraction (0.75) and the word size was optimized for each sample (**Supplementary Data 2**). The sequences of the resulting contigs were blasted (Blastx, E-value of ≤ 10E-6) against the Viridiplantae protein database (NCBI) and putative function annotated by Gene Onthology (cut-off = 55; GO weight = −5) using Blast2GO^33^. Filtered contigs for length (≥ 200 nt) and expression (TPKM > 1) were translated for the longest Open Reading Frame. Translated sequences longer than 20 amino acids, together with *Arabidopsis thaliana* protein sequences (TAIR10, www.arabidopsis.org) were analyzed by OrthoFinder^34^ to annotate Clusters of Orthologous Group (COG).

### Differential gene expression analysis

For each pair of species, read count estimation was carried out using CLC Genomics Workbench v9 software by mapping sequencing reads of each sample (default parameters, except similarity: 0.875 and length fraction: 0.75) to the transcriptome of the nickel hyperaccumulator species used as the reference. Statistical analyses to identify Differentially Expressed Genes (DEGs) were performed using the edgeR Bioconductor package^35^. To examine transcript abundance, reads per kilobase million (RPKM) were calculated. To evaluate the influence of the reference transcriptome, we additionally performed swapping analyzes using the transcriptome of the non-accumulator species as reference (**Supplementary Data 2**, **Supplementary Figure 3**).

### Molecular cloning

Predicted full-length coding region of *NcfiIREG2, GpruIREG1* and *LhavIREG2* were amplified from leaf cDNAs of the corresponding species, using high-fidelity Phusion polymerase (Thermo Scientific) with gene specific primers containing AttB recombination sequences (**Supplementary Data 11**). PCR products were first recombined into pDONOR207 (Invitrogen) and then in pDR195-GTW^36^ or pMDC83^37^ for expression in yeast and *N. caerulescens* respectively.

Artificial miRNA construct targeting *NcfiIREG2* was designed using the WMD3-Web microRNA Designer (http://wmd3.weigelworld.org). The amiRNA was engineered as previously described^38^ by PCR using the pRS300 backbone and specific primers (**Supplementary Data 11).** The *NcfiIREG2-*amiRNA precursor was recombined in pDONOR207 and then into the vector pK7GW2D^39^. All constructs were confirmed by restriction analysis and sequencing.

### Functional analyses of IREG/Ferroportin in yeast

IREG/ferroportin coding regions cloned into pDR195-GTW, as well as pDR195-AtIREG2^15^, were transformed by the lithium acetate method into the *Saccharomyces cerevisiae* BY4741 strain complemented by a functional *HIS3* gene. Yeast sensitivity to nickel was scored by yeast drop assay using serial dilution on histidine-free YNB agar medium containing 20 mM MES (pH5.5), supplemented or not with NiCl_2_. Empty pDR195 vector was used as control. Experiments were repeated twice with three independent transformants.

### Functional analysis of *NcfiIREG2* in transgenic plants

pK7GW2D-*NcfiIREG2-amiRNA* and *pMDC83-NcfiIREG2* were transformed into *N. caerulescens* subsp. *firmiensis* by *Rhizobium rhizogenes* (Arqua1 strain) mediated *in vitro* root transformation^40^. Transformed roots were selected using GFP fluorescence under a Leica MZ FLIII Fluorescence Stereo Microscope. Non-transformed roots were cut once a week until the whole root system was transgenic.

Independent lines transformed with pK7GW2D-*NcIREG2-amiRNA* were then transferred in hydroponic culture as described above for a week and then with the nutrient solution supplemented with 37.5 μM NiCl_2_ for 4 weeks. For each transgenic line, the root system was divided in two samples for RT-qPCR and elemental analyzes.

### Elemental analysis

Dry Environmental leaf samples were mineralized by HNO_3_. Multi-elemental analyzes were performed using ICP-AES (LAMA laboratory, IRD, New Caledonia) or MP-AES.

Transgenic root samples were washed twice with ice-cold 10 mM Na_2_EDTA and twice with ice-cold ultrapure water. Samples were dried at 65 ºC for 16 hours, weighted and then digested with 70 % HNO_3_ and H_2_O_2_ for a total of 8 h with temperature ramping from 80 to 120 °C. Elemental analyzes were performed using MP-AES (Agilent 4200, Agilent Technologies) and metal concentration was calculated by comparison with a metal standard solution.

### Confocal Imaging

Root transformed with pMDC83-*NcfiIREG2* were stained with 10 μg/ml propidium iodide (PI) and imaged on a Leica SP8X inverted confocal microscope

(IMAGERIE-Gif platform) with laser excitation at 488 nm and collection of emitted light at 495–550 nm for GFP and 600–650 nm for PI.

### Quantitative RT-PCR analyzes

Total RNA from *Ncfi* transgenic roots were extracted with TRI Reagent (Sigma-Aldrich). Total RNA from leaves of *Gpru* and *Ccap* were extracted with the CTAB-PVP method as described above. DNA was removed from all RNA samples by RNeasy Plant Mini kit on-column DNase I treatment. RNA (1 µg) were converted to cDNA by random priming using SuperScript III First-Strand reverse transcriptase (Invitrogen) according to the manufacturer’s instructions. Quantitative PCR analysis were performed on a LightCycler 96 using LightCycler 480 SYBR Green I Master Mix (Roche) with the following conditions: Initial denaturation (95 ºC, 300 s), followed by 40 cycles of amplification (95 °C, 15 s; 60 °C, 15 s; 72 °C, 15 s), and a melting curve (95 °C, 10 s; 65 ºC, 30 s and 95 ºC, 1s). Sequence information for Ccap (sample TIUZ) was obtained from the 1KP project^41^. Reference genes for *Ncfi* (*6-phosphogluconate decarboxylating 3, 6PGDC*, Ncfi_contig_3009; *Sister chromatid cohesion PDS5, PDS5*, Ncfi_contig_518), and *Gpru* and *Ccap* species (*Eukaryotic translation initiation factor 3, EIFL3*, Gpru_contig_12008; *Histone deacetylase 15, HDAC15*, Gpru_contig_14406) were selected from our RNA-Seq analyzes. Specific primers for *IREG/ferroportin* and reference genes were designed to produce amplicons of 100-200 nt (**Supplementary Data 11**). Relative gene expression was calculated using the primer efficiency correction method^42^. Two technical replicates and 3 independent biological replicates were used for each experiment.

### General statistics

Sample exploration and replication of sequencing data analysis were conducted using limma R package^43^, correlation analyses were conducted by linear regression using generic stats functions of R software^44^ and Heat-map clustering was carried out using heatmap.2 function of gplots R package^45^.

### Data availability

Raw sequence files, read count files and *de novo* assembled transcriptomes are available in the NCBI’s Gene Expression Omnibus under the SuperSeries accession number GSE116054 or SubSeries accession numbers GSE115411 (Brassicaceae), GSE116051 (Rubiaceae New Caledonia), GSE116050 (Rubiaceae Cuba), GSE116048 (Cunoniaceae), GSE116052 (Salicaceae) and GSE116049 (Euphorbiaceae).

